# Gluconeogenesis in the YSL-like tissue of cloudy catshark (*Scyliorhinus torazame*)

**DOI:** 10.1101/2024.03.29.587137

**Authors:** Marino Shimizu, Wataru Takagi, Yuki Sakai, Isana Kayanuma, Fumiya Furukawa

## Abstract

Glucose has important roles in the development of hematopoietic stem cells and brain of zebrafish, the vertebrate animal model; however, in most oviparous animals, the amount of maternally provided glucose in the yolk is scarce. For these reasons, developing animals need some ways to supplement glucose. Recently, it was found that developing zebrafish, a teleost fish, undergo gluconeogenesis in the yolk syncytial layer (YSL), an extraembryonic tissue that surrounds the yolk, utilizing yolk nutrients as substrates. However, teleost YSL is evolutionarily unique, and it is not clear how other vertebrates supplement glucose. In this study, we used cloudy catshark, an elasmobranch species which possesses a YSL-like tissue during development, and sought for possible gluconeogenic activities in this tissue. In the catshark yolk sac, an increase in glucose level was found, and our isotope tracking by ^13^C-labeled substrate combined with LC/MS analysis detected gluconeogenic activities with glycerol most preferred substrate. In addition, expression analysis for gluconeogenic genes showed that many of these were expressed at the YSL-like tissue, suggesting that cloudy catshark engages in gluconeogenesis in this tissue. The gluconeogenesis in teleost YSL and a similar tissue in elasmobranch species implies conservation of the mechanisms of yolk metabolism between these two lineages. Future studies on other vertebrate taxa will be helpful to understand the evolutionary changes in the modes of yolk metabolism that vertebrates have experienced.

## Introduction

Cartilaginous fishes branched from the bony vertebrates, including humans, around 450 million years ago, and did not experience the teleost-specific third-round whole genome duplication (1, 2). The reproduction of cartilaginous fishes exhibits highly diverse modes, ranging from oviparity to viviparity (3, 4). In oviparous species, the yolk serves as the sole energy source until the embryo begins feeding. However, it is not well understood how the yolk contents are utilized in the very early embryo where most organs are undeveloped. Recently, a novel metabolic phenomenon was found in developing zebrafish (*Danio rerio*), a teleost fish: yolk syncytial layer (YSL), an extraembryonic tissue surrounding the yolk, carries out gluconeogenesis using proteins and lipids, the main components of yolk (5). Gluconeogenesis is the series of enzymatic reactions where glucose is produced from non-sugar substrates such as amino acids and glycerol, and in vertebrate adult animals the liver and kidney are responsible for this function (6). Although glucose is considered to play important roles in normal development of brain and hematopoietic stem cells (7, 8), it is often very scarce in egg yolks (9), and the gluconeogenesis in YSL likely contributes to glucose supplementation in the teleost embryos. So far, YSL was believed to induce embryonic differentiation and regulate epiboly in the very early stages of development, and then contribute only to yolk transport and degradation. However, by the above finding, it was inferred that YSL is also responsible for metabolism of yolk components to actively synthesize and supplement the substances in need. Teleost YSL forms during the process of discoidal cleavage, a type of meroblastic cleavage (10), and it locates at the most inner layer of yolk sac membrane (YSM) after the end of epiboly. The existence of a tissue structurally similar to teleost YSL (YSL-like tissue) has been reported in the YSM of some elasmobranch species, which also undergo discoidal cleavage (11, 12), but its formation process is largely unknown. Thus, whether it has the same function as teleost YSL is unclear. Although the physiological role of YSL-like tissue in cartilaginous fish is unidentified, their yolk contains little glucose (13), similar to teleosts. Hence, we assumed that the YSL-like tissue is involved in the mechanism of yolk utilization to compensate for the lack of glucose.

In the present study, we used cloudy catshark (*Scyliorhinus torazame*), and examined changes in metabolite levels during their development. Because we confirmed increases in glucose levels in the yolk sac, we incubated the YSM with ^13^C-labeled metabolites and tracked their fate by liquid chromatography (LC) / mass spectrometry (MS) analysis. Gene expression analysis was also performed to further corroborate and localize the gluconeogenic activity. According to the results, we discuss possible metabolic roles of the YSL-like tissue in development of elasmobranchs and functional similarity to teleost YSL.

## Materials and Methods

### Animals

Cloudy catshark (*S. torazame*) were reared in 1000 or 3000-L tanks filled with recirculating seawater in the Atmosphere and Ocean Research Institute, the University of Tokyo. At this site, eggs were routinely laid by mature females and were reared in floating cages in 1000-L tanks under controlled light conditions (12 h light :12 h dark) at 16°C. The developmental stages (st.) of cloudy catshark embryos were identified using a detailed table for lesser spotted dogfish, *Scyliorhinus canicula* (14). Every effort was made to minimize the stress to the fish, and the embryos used were anesthetized by exposure to 0.05% tricaine before sampling. All experiments were approved by the Animal Ethics Committee of the Atmosphere and Ocean Research Institute of the University of Tokyo (P19-2). The present study was carried out in compliance with the ARRIVE guidelines.

### Metabolite analysis

Samples at st. 4, 24 (glucose only), 27, 31, and 32 were used to measure metabolites (n = 6). At st. 27, 31, and 32, embryo and yolk sac were separately analyzed. For every 1 g sample, 1 mL of distilled water, 2 mL 100% methanol, and 0.05 mL internal standard [L-methionine sulfone, 2-(N-morpholino) ethanesulfonic acid, D-camphor-10-sulfonic acid, 5 mM each] were added and homogenized. In addition, 2 mL chloroform was added and mixed. After keeping on ice for 10 min, the lysate was centrifuged at 13,000 *g* for 10 min at 4°C, and the aqueous phase was analyzed with an LC system (Prominence; Shimadzu, Kyoto, Japan) connected to a TripleTOF 5600+ mass spectrometer (AB Sciex, Framingham, MA). Shodex HILICpak VG-50 2D (Showa Denko, Tokyo, Japan) was used as the analytical column, and the mobile phase was solvent A: acetonitrile and solvent B: 0.5% aqueous ammonia. The initial concentration was 8.8% B, and the gradient conditions were 8.8% B (8 min), 95% B (14 min), 95% B (17 min), 8.8% B (19 min), and 8.8% B (24 min). The flow rate of the mobile phase was 0.2 mL/min and the column oven temperature was 60°C. For the determination of phosphorylated sugars and amino acids, Shodex HILICpak VT-50 2D (Showa Denko) was used, with solvent A: acetonitrile and solvent B: 25 mM ammonium formate at the constant ratio of A : B = 20 : 80 as mobile phase. The flow rate of the mobile phase was 0.3 mL/min and the column oven temperature was 60°C. The concentration of each metabolite in the samples was determined with reference to parallel measurement of standard solutions of stepwise dilutions.

### Isotope tracking

After anesthetizing st. 31 embryos (n = 9), the yolk sac was separated from the embryos. From the yolk sac, the yolk was removed, and the yolk sac membrane (YSM) was cut into six pieces. Glycerol-^13^C3 (Taiyo Nippon Sanso, Tokyo, Japan), Malate-^13^C4 (Taiyo Nippon Sanso), Sodium L-Lactate-^13^C3 (Cambridge Isotope Laboratories, Tewksbury, MA), L-Alanine-^13^C3 (Merck, Burlington, MA), and Glutamate-^13^C5, ^15^N (Taiyo Nippon Sanso) were diluted to 5 mM with Ringer’s solution for sharks (55 mM NaCl, 6.6 mM KCl, 1.4 mM Na_2_HPO_4_, 2.4 mM Na_2_SO_4_, 424 mM Urea, 56 mM Trimethylamine oxide, 10 mM HEPES, 4 mM MgCl_2_, 6.6 mM CaCl_2_, 3.6 mM NaHCO_3_; pH7.55). The pieces of dissected YSM were incubated for 3 h in the Ringer’s solution with or without (as controls) the labeled tracers. Metabolites were then extracted in the same manner as described above and subjected to LC/MS. The levels of tracer-containing metabolites, or isotopologues, were measured by targeting metabolites with excess mass (M) and expressed as % M+0.

### Real-time quantitative PCR

The total RNA was extracted from the embryo and the yolk sac membrane (YSM) of catshark at st. 27, 31, and 32 with TRI reagent (Molecular Research Center, Cincinnati, OH), digested with DNase I (Roche Diagnostics, Basel, Switzerland), and reverse transcribed with ReverTra Ace qPCR RT Kit (Toyobo, Osaka, Japan). qPCR was performed with Thermal Cycler Dice Real-time System II (Takara Bio, Shiga, Japan) and Luna Universal qPCR Master Mix (New England Biolabs, Ipswich, MA). Targeted genes related to gluconeogenesis and glucose transporter were: glucose-6-phosphatase, *g6pc1/g6pc*.*3*; phosphoenolpyruvate carboxykinase, *pck1/pck2*; glycerol-3-phosphate dehydrogenase, *gpd1/gpd1c/gpd1l/gpd2*; fructose-1,6-bisphosphatase, *fbp1*/*fbp2*; lactate dehydrogenase, *ldha/ldhb/ldhd*; pyruvate carboxylase, *pc*; glycogen synthase, *gys1*/*gys2*; glycogen phosphorylase, *pygb/pygm*; solute carrier family, *slc2a1/slc2a1b/slc2a2/slc2a5/slc2a8/slc2a11a/slc2a11b*). The primers used are listed in Table1. The PCR was performed in parallel with plasmid standard solutions of known concentrations (10^1^ ∼10^8^ copies/ µL) containing the respective target gene fragments, and the gene expression levels of each sample were calculated based on this PCR reaction. Dissociation curve analysis was performed after completion of PCR to determine the specificity of the amplified product.

### RNA probe synthesis

For *in situ* hybridization analysis, DIG-labeled RNA probes were prepared. First, target cDNA fragments were amplified with the primers listed in Table 1 and ligated into pGEM T-Easy vector (Promega, Madison, WI). The cDNA fragments were PCR amplified with M13 primers and purified with the FastGene Gel/PCR Extraction Kit (Nippon Genetics, Tokyo, Japan). The purified cDNA fragments were transcribed *in vitro* using T7 or SP6 RNA polymerase (Roche Diagnostics) in the presence of digoxigenin (DIG)-UTP. The resulting cRNA probe was digested with DNase I and purified with the RNeasy MinElute cleanup kit (Qiagen, Hilden, Germany).

#### *In situ* hybridization

To examine the localization of target gene transcripts, *in situ* hybridization analysis was performed according to a previous study (15). The embryos (or livers) and YSM at st. 27, 31, and 32 were fixed in 4% paraformaldehyde (PFA) in phosphate-buffered saline prepared with diethylpyrocarbonate-treated water (PBS-DEPC) overnight at 4°C, embedded in paraffin, sectioned into 5-µm slices, and placed onto glass slides. After deparaffinization, sections were digested with 2 µg/mL proteinase K for 30 minutes and postfixed with 4% PFA in PBS-DEPC for 15 minutes. The samples were then acetylated with 100 mM triethanolamine (pH 8.0) containing 0.25% acetic anhydride (16). After prehybridization with a hybridization mixture [HM^+^; 50% formaldehyde, 5x saline-sodium citrate buffer (SSC), 0.01% Tween 20, 500 µg/mL yeast tRNA, and 50 µg/mL heparin], samples were hybridized with DIG-labeled RNA probe (65 ng/ HM^+^ 200 μL) at 60°C overnight. After washing excess probes with 0.2 x SSC at 60°C, the samples were blocked with 5% skim milk in PBS-DEPC for 1 hour at 4°C and incubated with anti-DIG antibody (Roche Diagnostics) diluted 1:10,000 in the blocking buffer for 1 hour at 4°C. The mRNA signals were visualized by BCIP/NBT reaction, and the micrographs were obtained with a microscope (BZ-710, Keyence, Osaka, Japan).

### Statistics

All numerical data are expressed as mean ± standard error. The significance of difference between the means of groups were tested by one-way analysis of variance (ANOVA) followed by Tukey’s and Dunnet’s *post-hoc* test for metabolite analysis and metabolite tracking experiment, respectively. The results of qPCR analysis were analyzed by two-way ANOVA followed by Tukey’s test. To standardize the variations of values among groups, the data were log-normalized before the tests.

## Results

### Metabolite analysis

LC/MS analysis revealed that the levels of many metabolites in the embryo and yolk sac of catshark are highly variable during development (Fig.1, S1). Glucose concentration in the yolk sac increased approximately 100-fold from the egg just after spawning (st. 4) to st. 27, and it sharply increased after st. 24. The levels of glycogen, the storage form of glucose, tended to increase from st. 4 to st. 24 and then decreased. In addition, most metabolites in embryos increased with development, especially after st. 31.

**Fig. 1.**
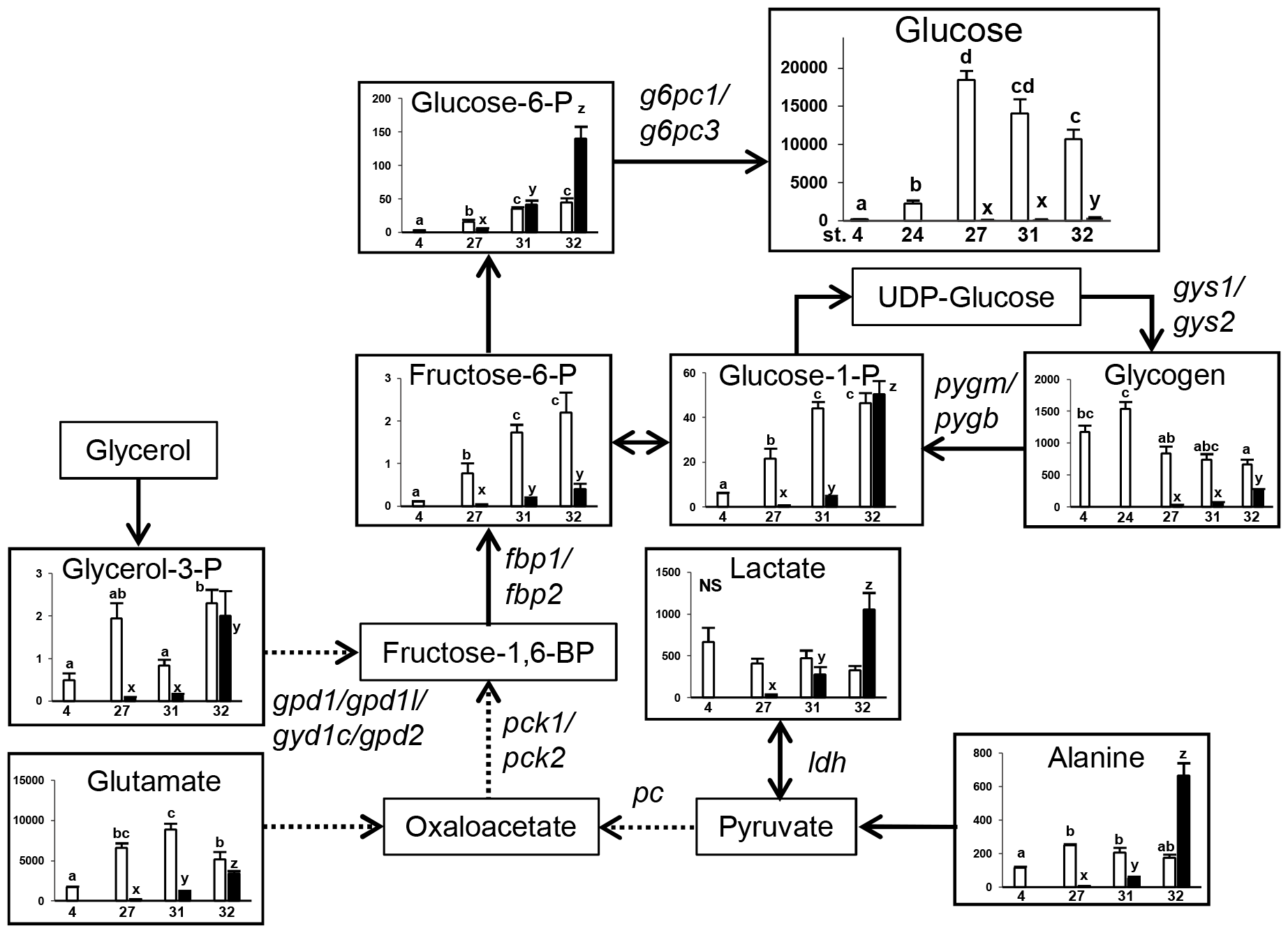
The metabolic pathway map showing changes in each metabolite levels per individual yolk sac (open column) or embryo (filled column) during development. The horizontal axes represent developmental stages {stages 4, 24 (glucose and glycogen only), 27, 31, 32} and the vertical axes represent nmol / sample. Data are presented as mean ± standard error (N = 6). Different letters indicate significant differences (*P* < 0.05) between groups. Tests for significant differences were performed by one-way ANOVA and Tukey’s *post-hoc* test separately for yolk sac or embryo samples after log transformation. Fructose-1,6BP, fructose 1,6-bisphosphate; -P, -phosphate. Genes responsible for respective pathways were shown beside the arrows. For the complete map with all metabolites measured in this study, please refer to Supplementary Figure S1.

### Isotope tracking

Isotope tracing was performed to determine the pathway by which glucose is produced (17). Glycerol, lactate, alanine, and glutamate were selected as possible substrates for gluconeogenesis, and the catshark YSM was incubated in these substrates labeled with ^13^C. As a result, levels of glucose with mass + 3 (M+3) were significantly elevated in the samples incubated with ^13^C-labeled glycerol and alanine (Fig. 2). Fructose-6-phosphate (F6P) M+3, glucose-6-phosphate (G6P) M+3, and glucose-1-phosphate (G1P) M+3, and sedoheptulose-7-phosphate (S7P), the intermediates of gluconeogenesis / glycogen metabolism / pentose phosphate pathway from the labeled substrates, were also significantly increased in samples incubated with labeled glycerol, lactate, and alanine (Fig. 2). The results for glucose M+3 in the samples which took labeled lactate was omitted due to interference of presumably other metabolites.

**Fig. 2.**
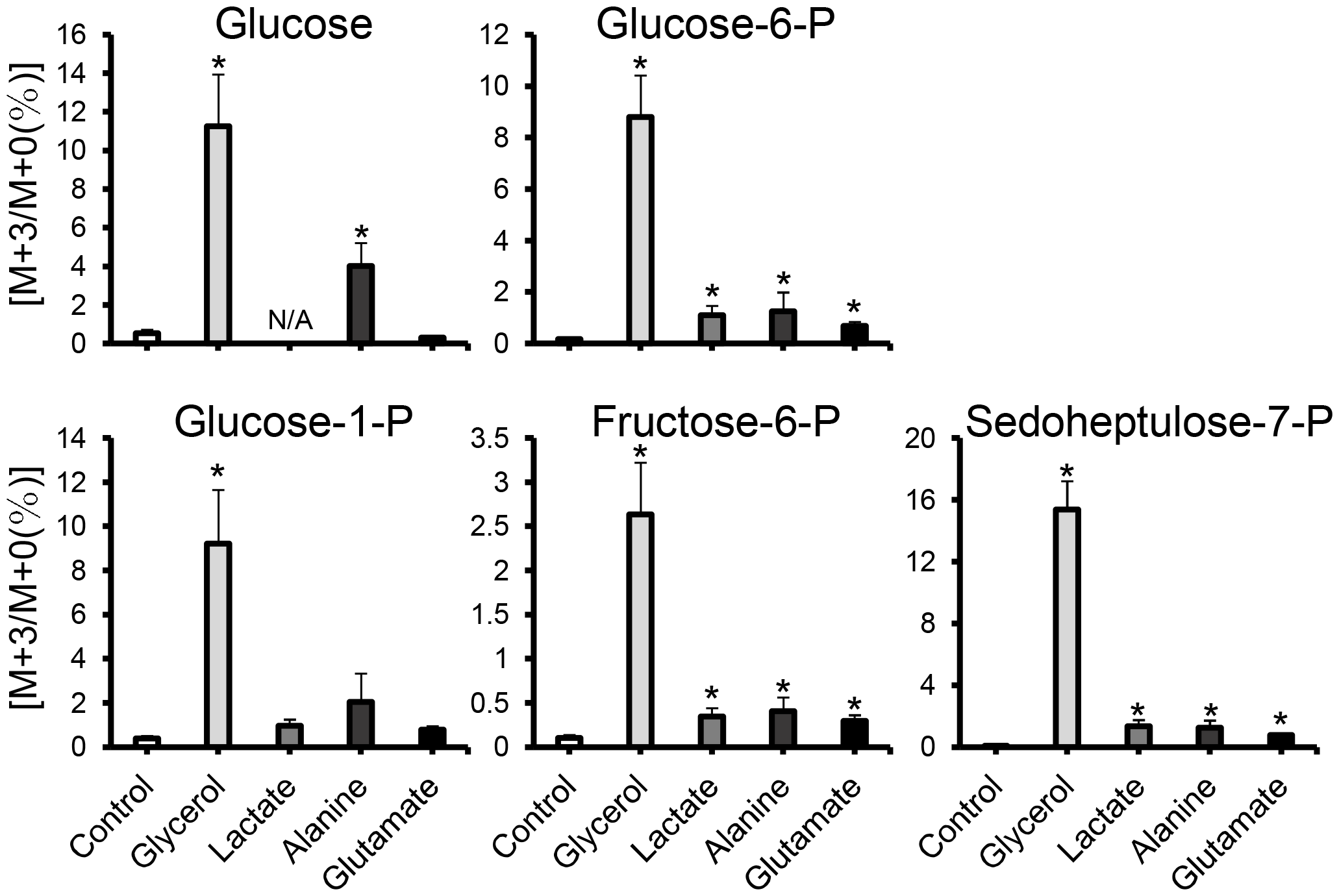
LC-MS-based isotope tracking of yolk sac membrane (YSM) in stage 31. Abundance of M+3 isotopologues of glucose, glucose-6-phosphate (-P), fructose-6-P, glucose-1-P, and sedoheptulose-7-P in the YSM after 3-hour incubation with ^13^C-labeled substrates. Horizontal axes show each ^13^C-labeled substrate and the control without them, and vertical axes show the levels of M+3 (as % M+0). Data are presented as mean ± standard error (N = 9), and asterisks (*) indicate significant differences (*P* < 0.05) between the control and each group. Tests for significant differences were conducted with one-way ANOVA followed by Dunnett’s test after log-transformation of the values. N/A, not available.

### Real-time PCR

Many of the gluconeogenesis-related genes showed differences in expression levels between embryonic body vs. yolk sac and among developmental stages (Fig. 3, S2). Among them, *g6pc1, fbp1, pc, pck1*, and *gys2* were highly expressed in the yolk sac and tended to be most highly expressed at st. 31. In the *slc2* glucose transporter gene family, only *slc2a2* was highly expressed in the yolk sac and at st.31.

**Fig. 3.**
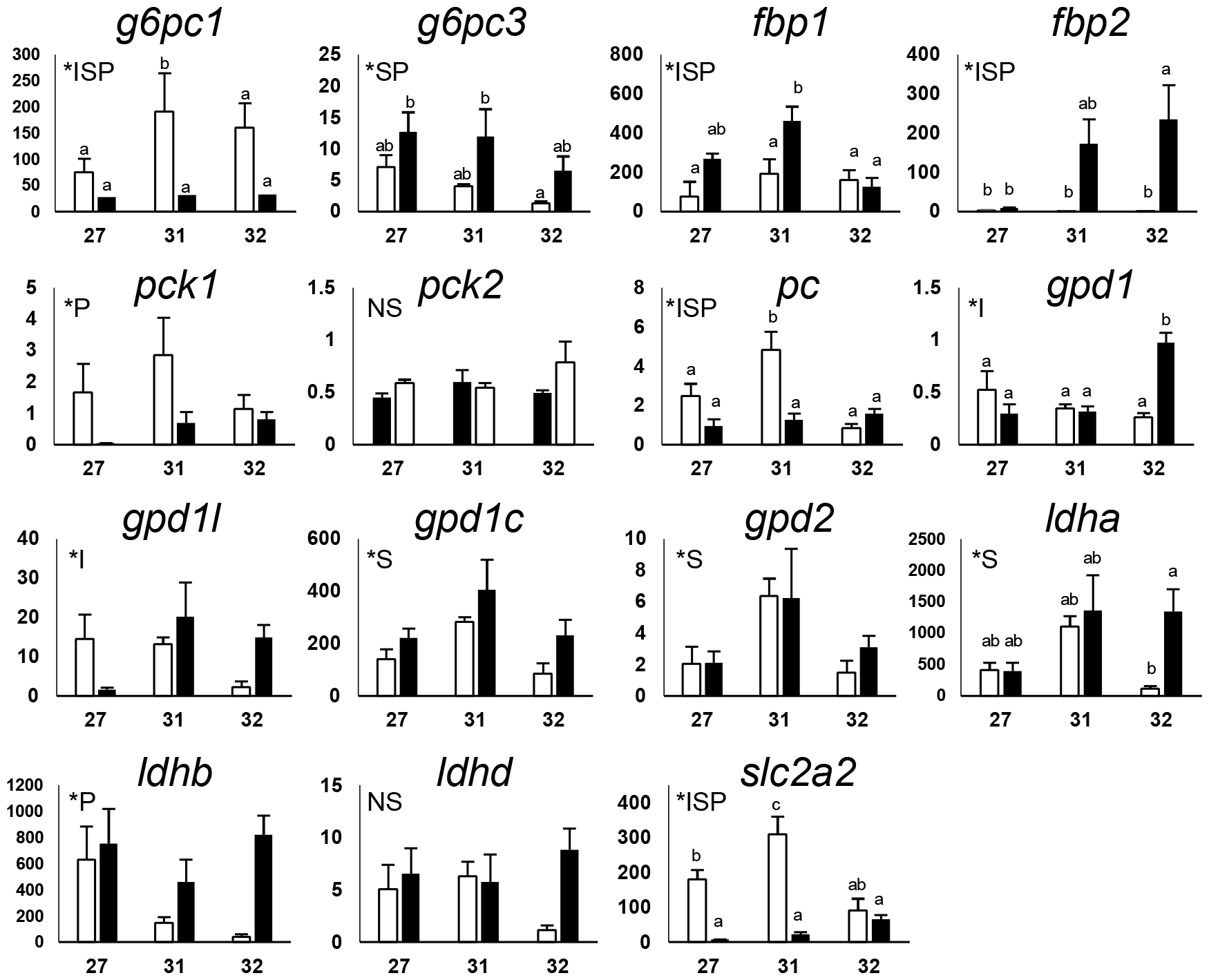
Changes in expression levels of the selected genes during development. The expression levels of gluconeogenesis-related genes and glucose transporter (GLUT) gene *slc2a2* were measured. Horizontal and vertical axes indicate developmental stages and mRNA levels (x 10^9^ copies / g RNA), respectively. Open and filled columns indicate the mRNA levels in the yolk sac membrane and embryos, respectively. Data are presented as mean ± standard error (N = 6), and different letters indicate significant differences (*P* < 0.05) between groups. Tests for significance were performed by two-way ANOVA and Tukey’s *post-hoc* test. *I indicates a significant interaction between the two factors (developmental stage and site), and *S and *P indicate significant main effects of developmental stage and the position (yolk sac or embryo), respectively. For other genes assessed in this study, please refer to Supplementary Figure S2.

#### *In situ* hybridization

For the mRNAs detected by real-time PCR, their localization was assessed in YSM and the embryonic livers at st. 27, 31, 32 by *in situ* hybridization. In the yolk sac membrane at st. 27, *g6pc1, fbp1, pck2, gpd1*, and *ldha* were expressed in YSL-like tissues and their nuclei (Fig. 4). In the same tissues, *pck1, gpd1l*, and *pc* were additionally expressed in st. 31 (Fig. 5). In st. 32, all transcripts assessed except *pck1* were found in YSL-like tissues and yolk sac endoderm (YSE), although the signals of *g6pc1* and *pc* were weak (Fig. 6). *pck1* was sporadically expressed in YSE at similar intervals. In the liver, transcripts of *pck1/pck2, gpd1/gpd1l, pc*, and *ldha* were expressed in st. 27 (Fig. S3). In addition to these genes, *g6pc1* and *fbp1* were expressed in st. 31 and 32 (Fig. S4, 5). In st. 31, the signals of *g6pc1* were sparse, probably in cells near the sinusoids (Fig. S4). These specific mRNA signals generated by antisense probes were not observed in the negative control slides subjected to sense probes.

**Fig. 4.**
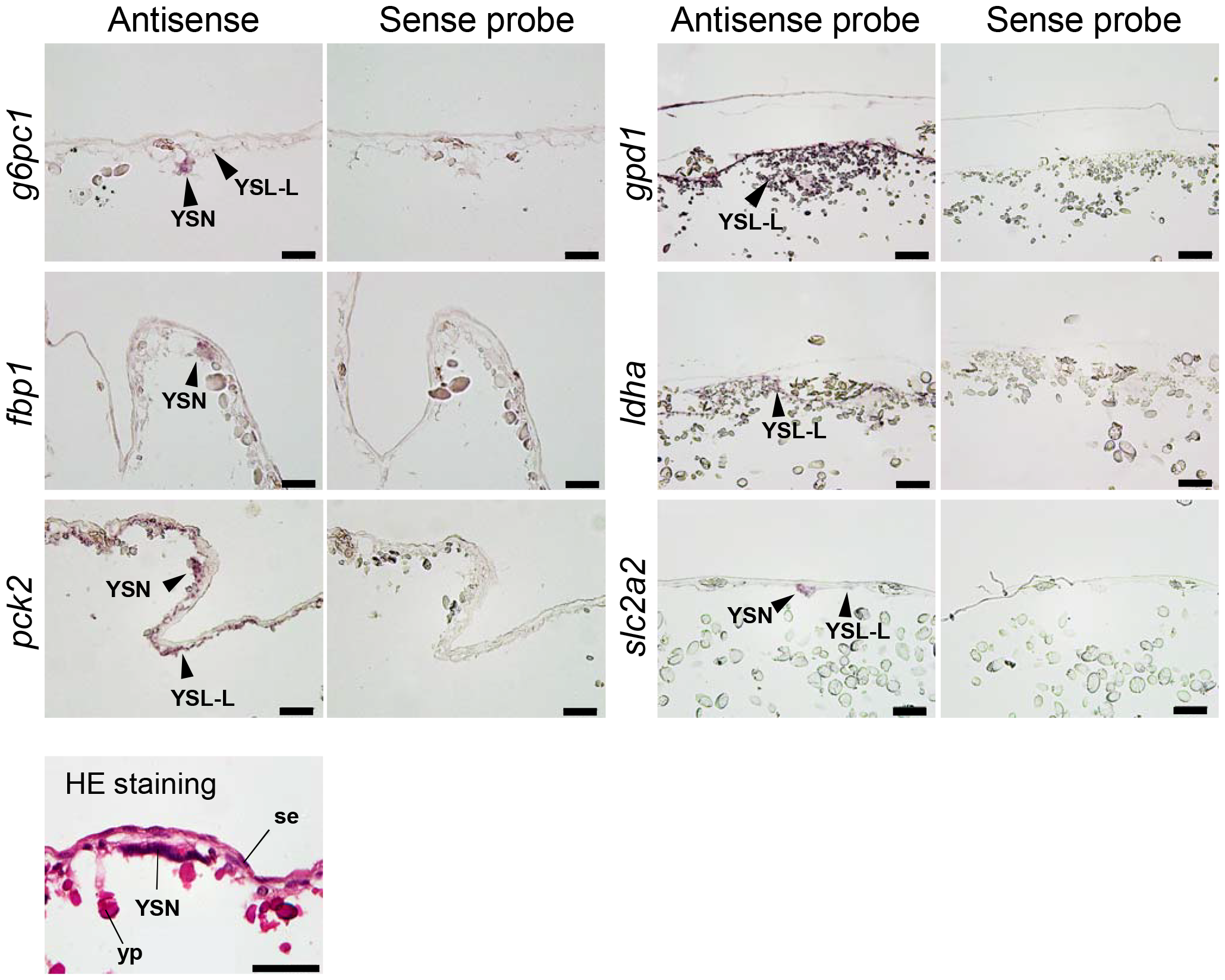
*In situ* hybridization analysis for *g6pc1, fbp1, pck2, gpd1, ldha*, and *slc2a2* at yolk sac membrane at stage 27. For all transcripts, sense probes (right) were used as negative controls for antisense probes (left), which showed true signals. Positive signals for transcripts are indicated by arrowheads. Hematoxylin and eosin (HE) staining shows general morphology of the yolk sac membrane in stage se, squamous epithelium; YSN, yolk syncytial nuclei; yp, yolk platelet. Scale bars = 50 μm.

**Fig. 5.**
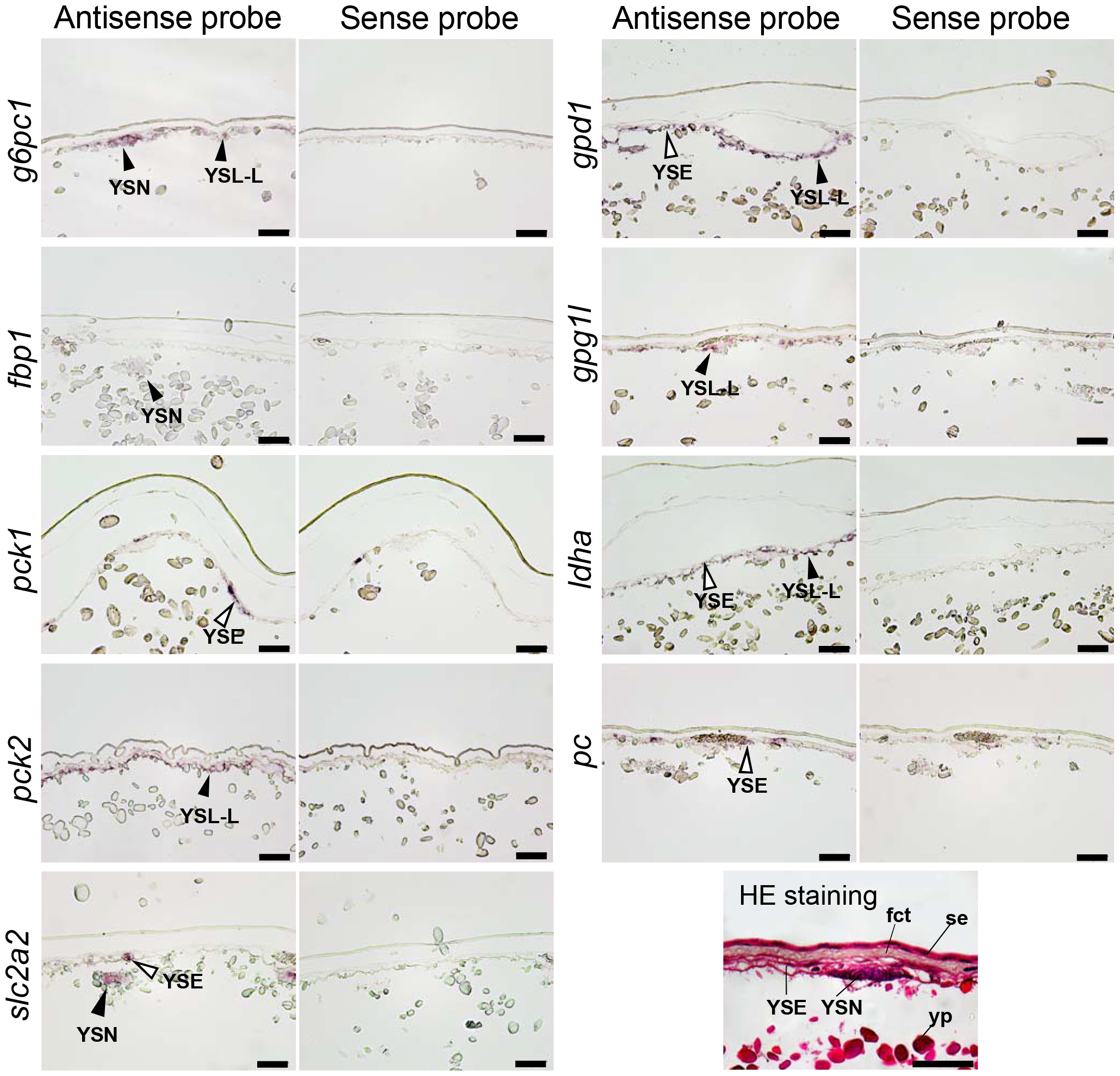
*In situ* hybridization analysis for *g6pc1, fbp1, pck1, pck2, gpd1, gpd1l, ldha, pc*, and *slc2a2* at yolk sac membrane at stage 31. Black arrowheads indicate signals in YSL-like tissue (YSL-L) and their nuclei (YSN), while white arrowheads indicate the signals in yolk-sac endoderm (YSE). Hematoxylin and eosin (HE) staining shows general morphology of yolk sac membrane in stage 31. se, squamous epithelium; fct, fibrillar connective tissue; yp, yolk platelet. Scale bars = 50 μm.

**Fig. 6.**
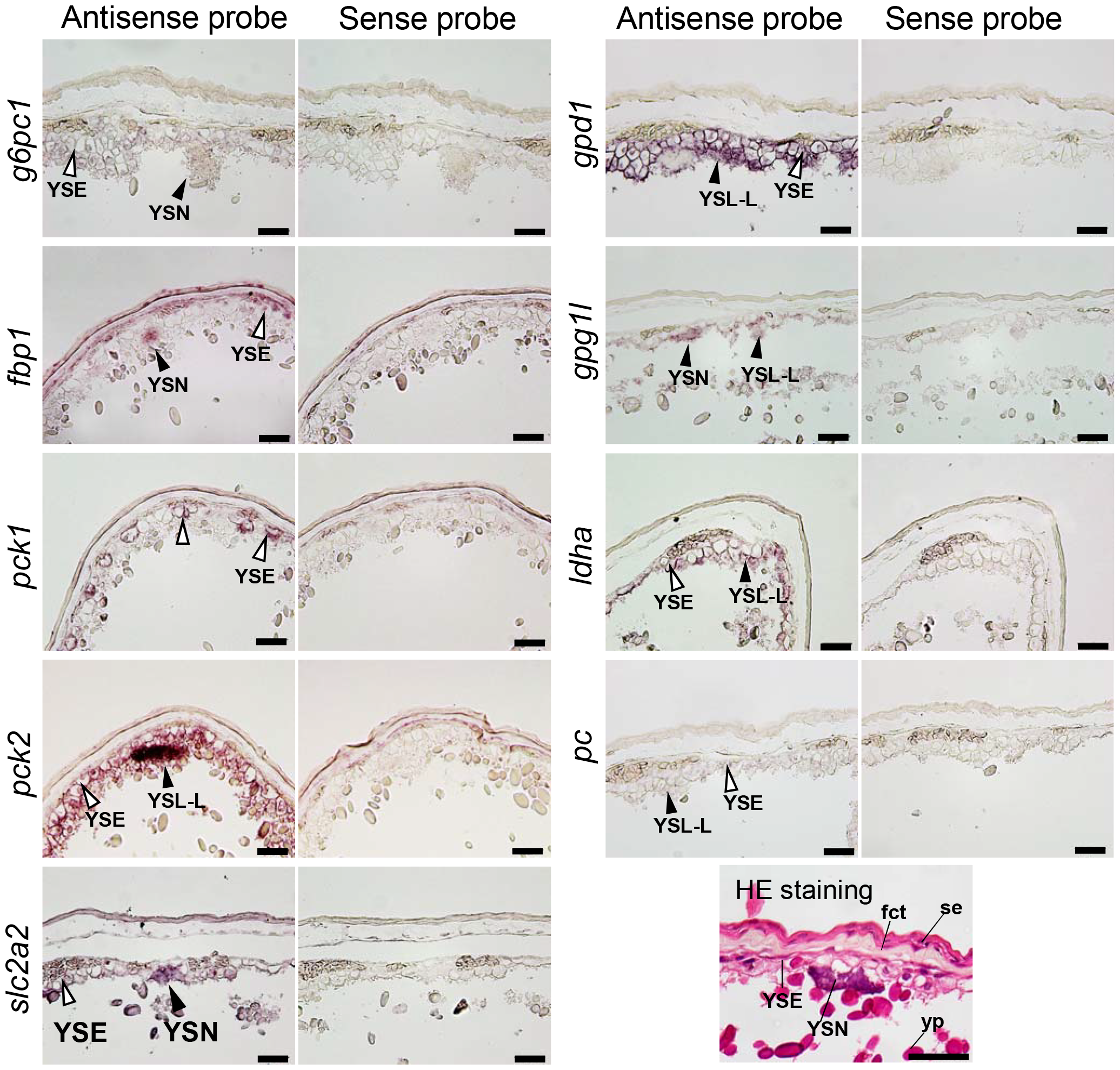
*In situ* hybridization for *g6pc1, fbp1, pck1, pck2, gpd1, gpd1l, ldha, pc*, and *slc2a2* at yolk sac membrane in stage 32. Black arrowheads indicate signals in YSL-like tissue (YSL-L) and their nuclei (YSN), while white arrowheads indicate the signals in yolk-sac endoderm (YSE). Hematoxylin and eosin (HE) staining shows general morphology of yolk sac membrane in stage 32. se, squamous epithelium; fct, fibrillar connective tissue; yp, yolk platelet. Scale bars = 50 μm.

## Discussion

Our metabolite analysis showed that catshark eggs just after spawning contain very little glucose, but it eventually increased, indicating that glucose is synthesized during the development (Fig. 1). In the lesser spotted dogfish, the development from fertilization to hatching is divided into 34 different stages according to the morphological characteristics of the embryo and yolk sac (14). In the present study, glucose content increased as the development proceeded, especially after st. 24 (Fig. 1). These changes occurred in the yolk sac samples, implying that YSM is responsible for the increase in glucose content during this period. Although glycogen was degraded in the yolk sac after st. 27, the decrease seemed inadequate to support the glucose production at this time, which led us to assess possible gluconeogenic activity in the YSM. Metabolite tracking in YSM incubated with ^13^C-labeled alanine, lactate, or glycerol revealed increases in M+3 isotopologues of F6P, G6P, G1P, and S7P, the intermediate metabolites for gluconeogenesis, glycogen metabolism, and pentose phosphate pathway (Fig. 2). Also, glucose M+3 was elevated in YSM incubated with labeled alanine or glycerol. These M+3 isotopologues are most likely those containing three ^13^C, which were passed through the gluconeogenic pathway from the substrates added. These results suggest that the labeled substrates are used to produce glucose and glycogen in YSM of the catshark, which also supports the results of the metabolite analysis. Among the substrates added, glucose was most efficiently produced from glycerol. The yolk of lesser spotted dogfish contains about 20% lipid on dry weight basis (18), and the total dry mass of the yolk is reduced by about 20% by hatching (19). On the other hand, approximately 44% of yolk lipids alone are lost, suggesting that these lipids are actively used during development for lipid membrane, energy source, and/or metabolic substrate. Lipids such as triglycerides yield glycerol upon degradation, which may subsequently function as a substrate for gluconeogenesis. These previous studies support the results in the tracer experiments, and taken together, elasmobranchs likely use glycerol as a preferred substrate for gluconeogenesis during development.

In our qPCR analysis, many of the gluconeogenesis-related genes were expressed in the yolk sac (Fig. 3), and subsequent *in situ* hybridization analysis also supported these results. The catshark YSM consists of: (from the outside) two layers of squamous epithelium, fibrous connective tissue, basement membrane, vascular endothelium, YSE, and YSL-like tissue (11, 19, 20). Within YSM, many of the gluconeogenesis-related genes were expressed in the YSL-like tissue and/or YSE, even when the signal was absent in the embryonic liver, after st. 31 (Fig. 4, 5, 6). *pck1* was found only in a portion of YSE, implying that YSE consists of more than one type of cells. *g6pc1*, a gene required to produce glucose from G6P, showed a robust signal in YSL-like tissue at st. 31, while it became weak at st. 32 (Fig. 5, 6), supporting our qPCR data (Fig. 3). These results suggest the existence of active gluconeogenesis in the YSL-like tissue at st.31, and marginal gluconeogenic activity in YSM (YSL-like tissue and YSE) after st. 32. In catsharks, the anterior portion of the yolk capsule opens during st. 31, a phenomenon called pre-hatching, and at similar timing, yolk reaches the gut through the yolk stalk and starts to be absorbed (21). These events trigger increases in the expression of various genes responsible for amino acid transport, lipid absorption, and digestive enzymes in the gut and yolk sac (12). The results of this study suggest that the YSL-like tissue of catshark actively performs metabolism such as gluconeogenesis using the yolk before the development of many organs. Later, however, its metabolic functions are shifted to the embryo (liver, intestine, kidney, etc.), and YSL-like tissue may focus more on absorption and transport of the yolk together with the gut and YSE (Fig. 7). Similar phenomenon was observed in this species for urea synthesis, where this osmolyte was actively synthesized in YSM at early stages, but later the liver takes over this critical physiological function (22).

**Fig. 7.**
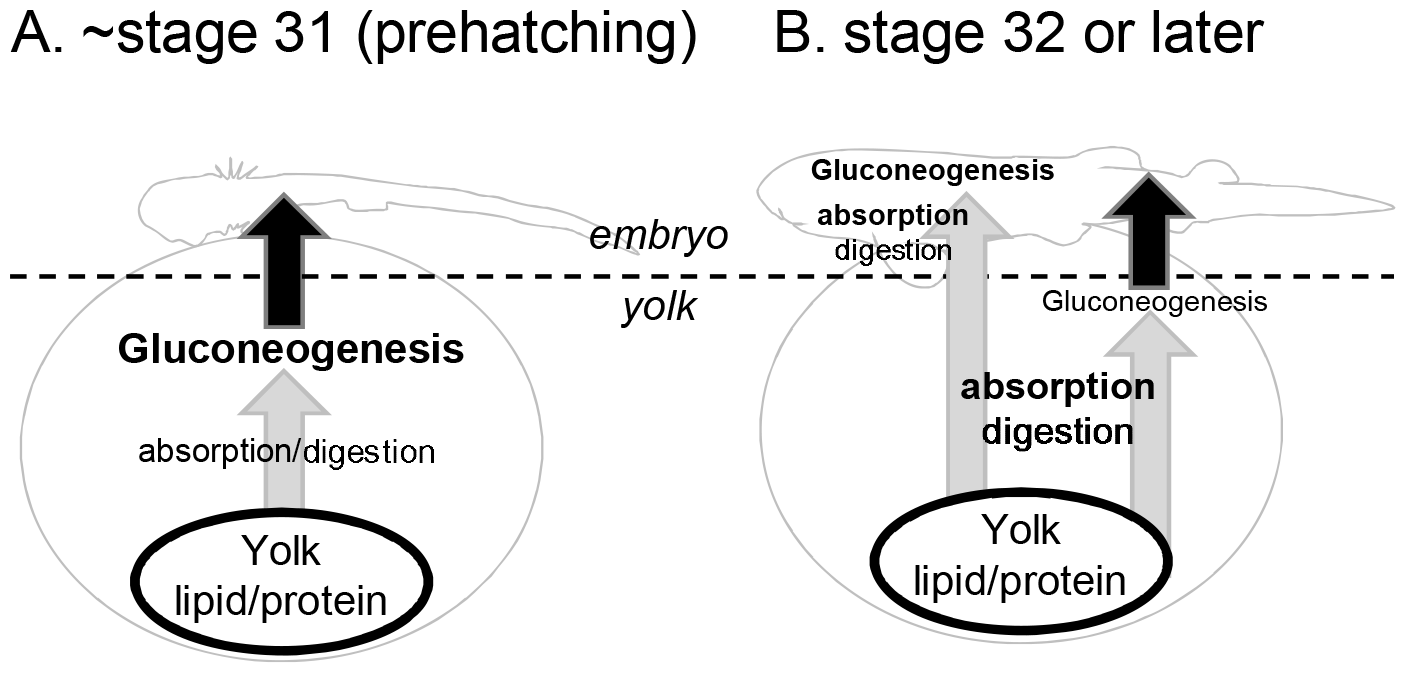
Model of functional changes in the yolk sac over the near-prehatching stages. A. Until stage 31, where pre-hatching takes place, yolk degradation, absorption, and gluconeogenesis occur mainly in the yolk sac; B. At stage 32 or later, yolk degradation and absorption become more active in both embryonic gut and yolk sac, and glucogenesis and other processes occur actively in the embryo.

In the cloudy catshark, many orthologous genes of those expressed in zebrafish YSL were also expressed in YSL-like tissues. The same was true for the *slc2* family genes encoding glucose transporters, where *slc2a2* was YSL-like type as in zebrafish (23). The functional similarity between zebrafish YSL and catshark YSL-like tissue in terms of gluconeogenesis may raise some questions: do these tissues share a common evolutionary origin, and are they the products of parallel evolution occurring on the same molecular basis (24)? However, reports exist that YSL in teleost fishes and shark YSL-like tissues have different nuclear origins. In teleosts, the YSL nuclei are formed by the disintegration of cells at the marginal blastomere (limbal cells) and their fusion with the cytoplasmic layer on the yolk surface (10). On the other hand, the elasmobranch YSL-like tissue has primary nuclei derived from the blastoderm and secondary nuclei from the endoderm (25). The primary nuclei are believed to be derived from those left behind in the yolk when the segmentation cavity is broken up into yolk and embryo, and it is not clear whether the disintegration of the blastodisc marginal cells occur. Thus, many aspects of the formation, structure, and function of the elasmobranch YSL-like tissues are unknown and may differ from those of teleost YSL. For now it is not possible to determine whether the YSL / YSL-like tissues in these two taxa are the products of parallel evolution or of convergent evolution, the latter occurring on a different molecular basis (26). To answer these questions, it is necessary to observe and analyze in detail the morphological and molecular aspects of the formation of YSL-like tissues, as well as tracking fates of counterpart cell lineages in other bony fishes and cyclostomes.

The present study showed that cloudy catsharks undergo gluconeogenesis using glycerol and other substrates in YSM, where the YSL-like tissue likely takes the central role. Gluconeogenesis before organ development is essential and may be a widely conserved phenomenon in many vertebrate animals. It is still unclear whether it was acquired in the common ancestor and modified during evolution to suit each taxon, or whether it was acquired independently. Future studies of the sites of gluconeogenesis in other vertebrate taxa, together with the YSL-like tissue of catshark, will help us trace the roots of this tissue and to elucidate early developmental changes that vertebrates experienced during their evolution.

## Acknowledgement

We thank all the staff of Oarai Aquarium for generously providing us the adult cloudy catshark. Prof. Susumu Hyodo (Laboratory of Physiology, Atmosphere and Ocean Research Institute, The University of Tokyo) helped us conduct the experiments and allowing use of lab equipments. This work was partially supported by Grant-in-Aid for Young Scientists [no.18K14524], Grant-in-Aid for Exploratory Research [no. 21K19276], and Grant-in-Aid for scientific Research (B) [no. 22H02426] from JSPS, and Kitasato University Research Grant for Young Researchers to Fumiya Furukawa.

---

Table1. List of primers used in this study

ISH; in situ hybridization, qPCR; quantitative PCR (Real-time PCR)

