## Supplemental Fig. 1-5 for "Gluconeogenesis in the YSL-like tissue of cloudy catshark (*Scyliorhinus torazame*)"

### Slide 1
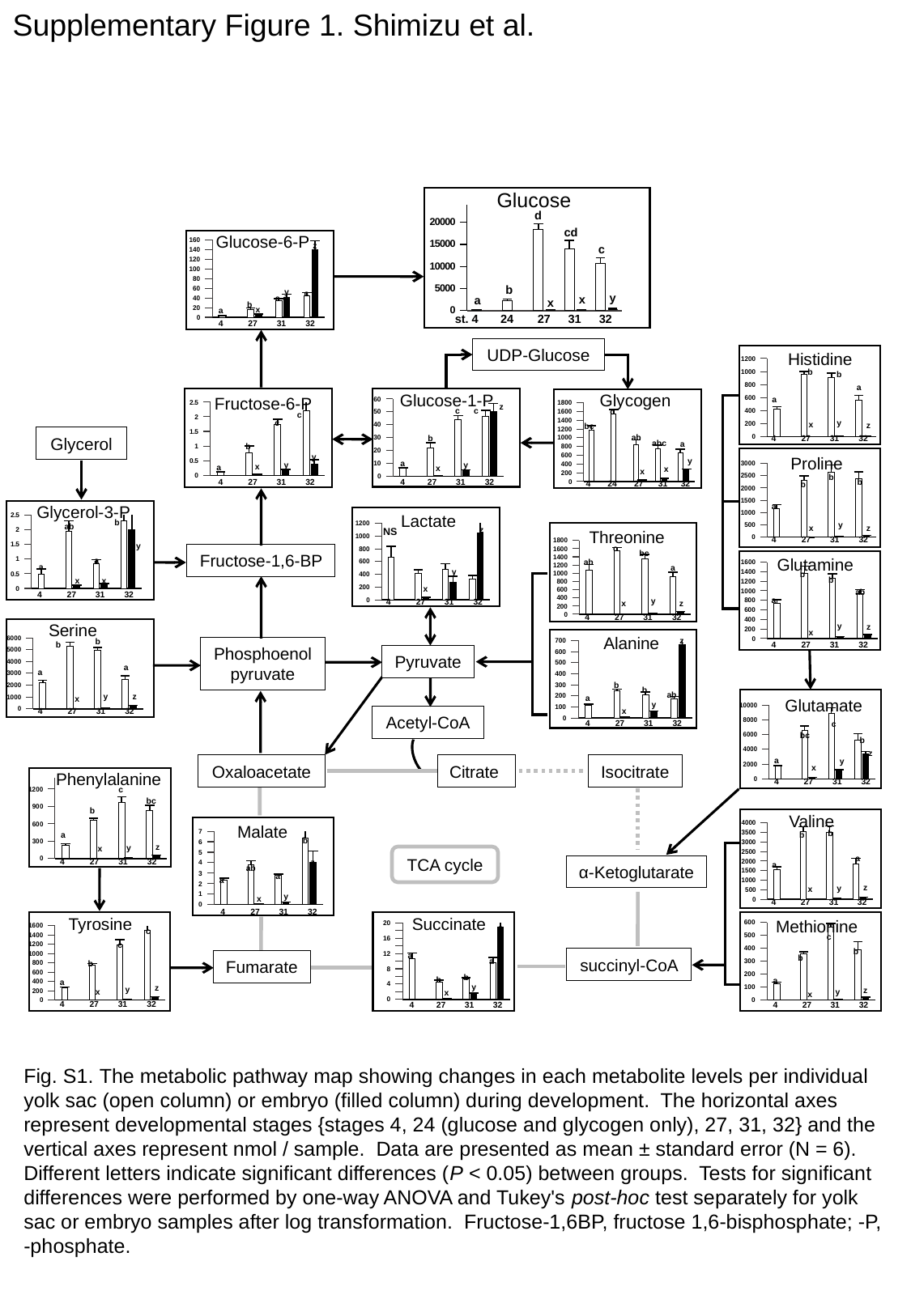

Supplementary Figure 1. Shimizu et al.
Glucose
#### Chart
| Category | 卵黄 | 胚 |
|---|---|---|
| 4 | 167.74964017596218 | None |
| 24.5 | 2251.444588985011 | None |
| 27 | 18470.129667056437 | 93.95654709071289 |
| 31 | 14074.21544794358 | 152.50105580195557 |
| 32 | 10688.383832708056 | 417.01000271167345 |d
cd
Glucose-6-P
#### Chart
| Category | 卵黄 | 胚 |
|---|---|---|
| 4 | 2.6341228865570065 | None |
| 27 | 15.71938774646644 | 5.708048788662608 |
| 31 | 34.776776974384305 | 40.9614808574786 |
| 32 | 44.71395267262625 | 139.97833889453776 |z
c
b
y
c
y
x
c
a
x
b
x
a
st. 4
24
27
31
32
4
27
31
32
UDP-Glucose
Histidine
#### Chart
| Category | 卵黄 | 胚 |
|---|---|---|
| 4 | 424.031538942051 | None |
| 27 | 954.8233673231172 | 0.7509804406176456 |
| 31 | 901.4466781292971 | 1.9410682893529634 |
| 32 | 562.1090464480056 | 11.740717561802654 |b
b
a
Glycogen
Glucose-1-P
Fructose-6-P
a
#### Chart
| Category | 卵黄 | 胚 |
|---|---|---|
| 4 | 6.136171636167132 | None |
| 27 | 21.68915331337134 | 0.6597870368199363 |
| 31 | 44.15342074958792 | 4.217949799614286 |
| 32 | 46.48495509308098 | 50.39146595499741 |
#### Chart
| Category | 卵黄 | 胚 |
|---|---|---|
| 4 | 0.10573481434313657 | None |
| 27 | 0.7668931922055998 | 0.03313586619389808 |
| 31 | 1.7254257329973461 | 0.16247737450524896 |
| 32 | 2.202289198861871 | 0.396806616072299 |
#### Chart
| Category | 卵黄嚢 | 胚 |
|---|---|---|
| 4 | 1177.9297376562008 | None |
| 24 | 1536.5034190425347 | None |
| 27 | 835.0529408530256 | 29.51191689192234 |
| 31 | 741.3955465344551 | 56.999565165550145 |
| 32 | 662.5854456428716 | 260.08951666730496 |
z
c
c
c
c
y
c
x
z
bc
ab
b
Glycerol
4
27
31
32
abc
a
b
y
Proline
y
a
y
y
x
#### Chart
| Category | 卵黄 | 胚 |
|---|---|---|
| 4 | 1158.7480706975784 | None |
| 27 | 2296.9701812644935 | 2.424814845195574 |
| 31 | 2621.1441310307832 | 13.581774714077865 |
| 32 | 2379.370694341848 | 72.90737527879143 |a
x
x
x
b
4
27
31
32
b
4
27
31
32
4
27
31
32
24
b
a
Glycerol-3-P
Lactate
#### Chart
| Category | 卵黄 | 胚 |
|---|---|---|
| 4 | 0.49643855439979984 | None |
| 27 | 1.939898657579591 | 0.0799527242950941 |
| 31 | 0.8365060365851059 | 0.14781447113463844 |
| 32 | 2.3002067676153644 | 1.9993464070334195 |b
y
ab
#### Chart
| Category | 卵黄 | 胚 |
|---|---|---|
| 4 | 663.9223540267511 | None |
| 27 | 409.79689217619426 | 32.55455911656072 |
| 31 | 470.6237919687945 | 278.39758789773055 |
| 32 | 329.1926404529152 | 1051.1101660688514 |z
x
z
NS
Threonine
4
27
31
32
#### Chart
| Category | 卵黄 | 胚 |
|---|---|---|
| 4 | 1078.9642784001467 | None |
| 27 | 1541.224229712336 | 4.233686582279698 |
| 31 | 1342.2568841506018 | 15.559740289750083 |
| 32 | 919.5735987754069 | 61.779023993734484 |y
c
bc
Fructose-1,6-BP
Glutamine
a
ab
#### Chart
| Category | 卵黄 | 胚 |
|---|---|---|
| 4 | 726.8481638501642 | None |
| 27 | 1356.9248958280307 | 2.041844534515129 |
| 31 | 1259.9603421424993 | 32.28967033258512 |
| 32 | 932.0874814649611 | 71.67512113048866 |a
a
y
b
b
x
x
x
ab
4
27
31
32
a
y
4
27
31
32
z
x
4
27
31
32
Serine
y
z
x
Alanine
z
b
#### Chart
| Category | 卵黄 | 胚 |
|---|---|---|
| 4 | 2206.870278296494 | None |
| 27 | 5296.430498937341 | 29.13632644269345 |
| 31 | 4964.705073288245 | 82.77211944317217 |
| 32 | 2501.470427720514 | 276.9175829312562 |
#### Chart
| Category | 卵黄 | 胚 |
|---|---|---|
| 4 | 116.70702786249572 | None |
| 27 | 249.1313341259611 | 5.818889389733028 |
| 31 | 206.77596233863048 | 54.5781723092885 |
| 32 | 174.5308933413588 | 666.3277882740393 |4
27
31
32
b
Phosphoenolpyruvate
Pyruvate
a
a
b
b
ab
z
y
a
x
Glutamate
y
#### Chart
| Category | 卵黄 | 胚 |
|---|---|---|
| 4 | 1709.7922291991501 | None |
| 27 | 6576.187204594059 | 196.19054179673333 |
| 31 | 8850.090730465488 | 1132.8983622969058 |
| 32 | 5197.805543501571 | 3402.4736555482373 |x
4
27
31
32
Acetyl-CoA
4
27
31
32
c
bc
b
z
a
y
Oxaloacetate
Citrate
Isocitrate
x
Phenylalanine
#### Chart
| Category | 卵黄 | 胚 |
|---|---|---|
| 4 | 229.24728208114698 | None |
| 27 | 655.5446464663734 | 3.048797153666742 |
| 31 | 965.0422961740184 | 11.261178292733051 |
| 32 | 823.3449313424549 | 52.754656273390744 |4
27
31
32
c
bc
b
Valine
Malate
#### Chart
| Category | 卵黄 | 胚 |
|---|---|---|
| 4 | 1564.3395287403728 | None |
| 27 | 3552.2517026607857 | 10.188957805827384 |
| 31 | 3509.689541355589 | 33.49497394824317 |
| 32 | 1852.7291589412246 | 85.29926699348867 |
b
a
b
#### Chart
| Category | 卵黄 | 胚 |
|---|---|---|
| 4 | 2.3017670171348548 | None |
| 27 | 3.824933849312151 | 0.04083582774385672 |
| 31 | 2.761876417284066 | 0.21926541653298393 |
| 32 | 6.392824470592563 | 3.996240585853441 |b
z
y
x
TCA cycle
a
4
27
31
32
z
a
α-Ketoglutarate
ab
a
a
z
y
x
y
x
4
27
31
32
4
27
31
32
Succinate
Tyrosine
Methionine
#### Chart
| Category | 卵黄 | 胚 |
|---|---|---|
| 4 | 125.18664830138242 | None |
| 27 | 359.40116745116757 | 2.207587329299345 |
| 31 | 562.1075239665442 | 5.583366738022594 |
| 32 | 390.6880890844145 | 19.51838350588326 |
#### Chart
| Category | 卵黄 | 胚 |
|---|---|---|
| 4 | 10.67875710405337 | None |
| 27 | 4.522313686447544 | 0.16326122830296994 |
| 31 | 5.275972442882227 | 1.334112826567635 |
| 32 | 9.618112734143219 | 18.839763347123917 |z
#### Chart
| Category | 卵黄 | 胚 |
|---|---|---|
| 4 | 257.9022104930909 | None |
| 27 | 750.2147972852936 | 3.214699425548569 |
| 31 | 1178.880034090801 | 11.002274276769656 |
| 32 | 1503.4309459913075 | 63.25425582798832 |c
c
c
b
a
b
succinyl-CoA
a
Fumarate
b
b
b
a
a
y
z
y
z
x
y
x
x
4
27
31
32
4
27
31
32
4
27
31
32
Fig. S1. The metabolic pathway map showing changes in each metabolite levels per individual yolk sac (open column) or embryo (filled column) during development. The horizontal axes represent developmental stages {stages 4, 24 (glucose and glycogen only), 27, 31, 32} and the vertical axes represent nmol / sample. Data are presented as mean ± standard error (N = 6). Different letters indicate significant differences (P < 0.05) between groups. Tests for significant differences were performed by one-way ANOVA and Tukey's post-hoc test separately for yolk sac or embryo samples after log transformation. Fructose-1,6BP, fructose 1,6-bisphosphate; -P, -phosphate.

### Slide 2
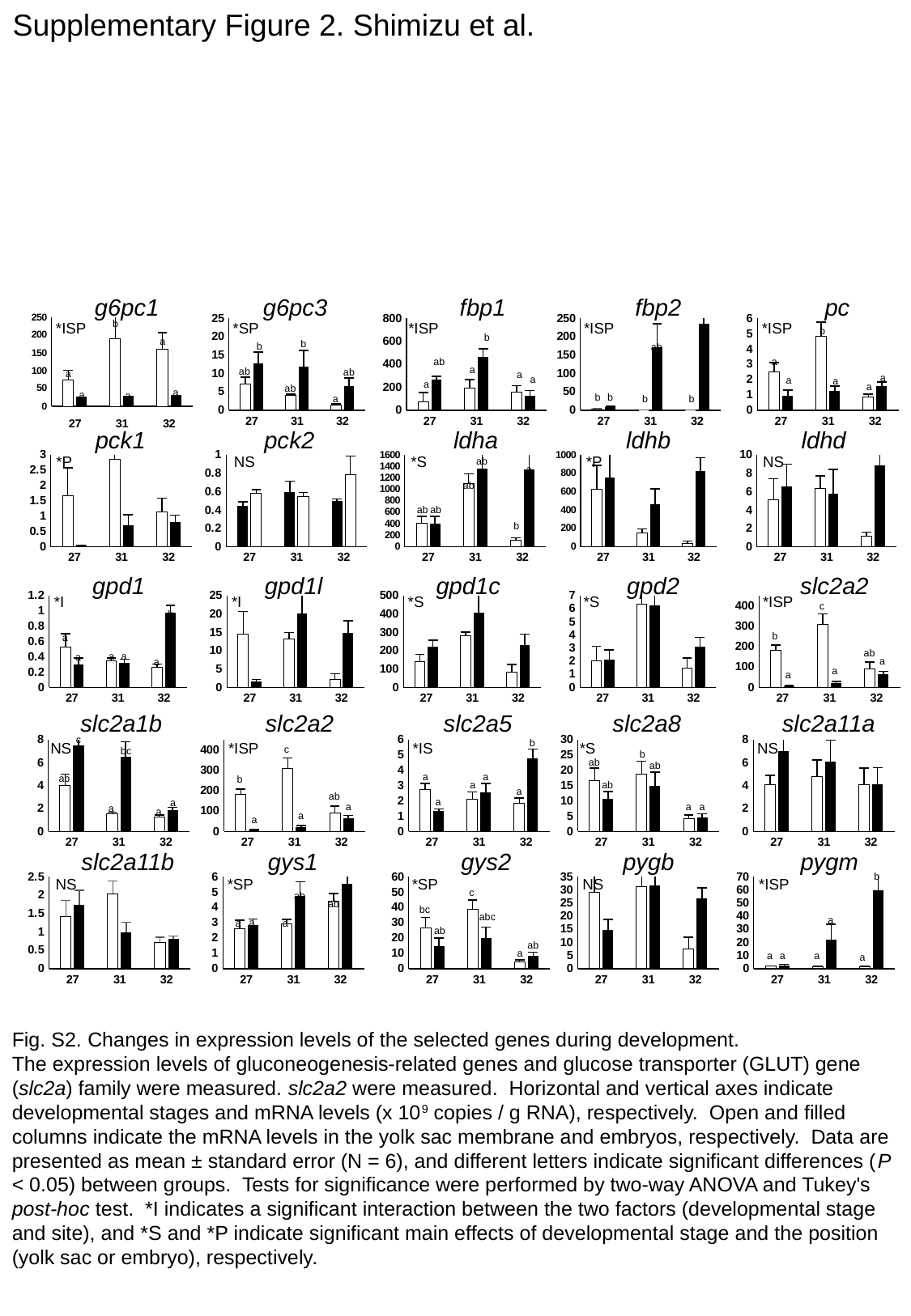

Supplementary Figure 2. Shimizu et al.
g6pc1
g6pc3
fbp1
fbp2
pc
#### Chart
| Category | embryo | |
|---|---|---|
| 27 | 74.68938888888889 | 27.0 |
| 31 | 190.76577777777777 | 31.0 |
| 32 | 160.20666666666668 | 32.0 |
#### Chart
| Category | 卵黄嚢 | 胚 |
|---|---|---|
| 27 | 7.07488888888889 | 12.686277777777777 |
| 31 | 4.000444444444445 | 11.93177777777778 |
| 32 | 1.2754666666666665 | 6.536499999999999 |
#### Chart
| Category | embryo | yolk |
|---|---|---|
| 27 | 74.68938888888889 | 269.16013333333336 |
| 31 | 190.76577777777777 | 461.2283888888889 |
| 32 | 160.20666666666668 | 125.20975000000001 |
#### Chart
| Category | 卵黄嚢 | 胚 |
|---|---|---|
| 27 | 2.249055555555555 | 9.196388888888889 |
| 31 | 0.6178333333333333 | 172.34827777777778 |
| 32 | 0.29341666666666666 | 234.8799444444444 |
#### Chart
| Category | 卵黄嚢 | 胚 |
|---|---|---|
| 27 | 2.4770000000000003 | 0.9441666666666667 |
| 31 | 4.823166666666666 | 1.2448666666666668 |
| 32 | 0.8432000000000001 | 1.5627777777777778 |b
*ISP
*SP
*ISP
*ISP
*ISP
a
b
b
a
b
b
ab
ab
a
a
ab
ab
a
a
a
a
a
a
a
a
ab
a
a
a
b
b
b
b
a
pck1
pck2
ldha
ldhb
ldhd
#### Chart
| Category | yolk | embryo |
|---|---|---|
| 27 | 1.6623333333333334 | 0.03533333333333333 |
| 31 | 2.8457777777777777 | 0.6926111111111112 |
| 32 | 1.1270833333333334 | 0.8032777777777778 |
#### Chart
| Category | embryo | yolk |
|---|---|---|
| 27 | 0.4492777777777777 | 0.5850000000000001 |
| 31 | 0.5954444444444444 | 0.5429444444444445 |
| 32 | 0.4953888888888888 | 0.7847333333333333 |
#### Chart
| Category | yolk | embryo |
|---|---|---|
| 27 | 407.6332777777779 | 395.2665555555555 |
| 31 | 1103.7047222222222 | 1357.9264666666666 |
| 32 | 110.60558333333333 | 1344.1897222222221 |
#### Chart
| Category | yolk | embryo |
|---|---|---|
| 27 | 630.4171666666667 | 754.5000666666666 |
| 31 | 147.053 | 459.61033333333336 |
| 32 | 39.036 | 822.8364444444445 |
#### Chart
| Category | yolk | embryo |
|---|---|---|
| 27 | 5.070944444444444 | 6.5631666666666675 |
| 31 | 6.309611111111111 | 5.780666666666667 |
| 32 | 1.1528333333333334 | 8.828733333333334 |*P
NS
*S
*P
NS
ab
a
ab
ab
ab
b
#### Chart
| Category | yolk | embryo |
|---|---|---|
| 27 | 0.5225555555555556 | 0.2971666666666667 |
| 31 | 0.3451666666666667 | 0.3163888888888889 |
| 32 | 0.26216666666666666 | 0.9756111111111109 |
#### Chart
| Category | yolk | embryo |
|---|---|---|
| 27 | 14.52961111111111 | 1.5733333333333335 |
| 31 | 13.124055555555556 | 20.1494 |
| 32 | 2.1906666666666665 | 14.882666666666667 |
#### Chart
| Category | yolk | embryo |
|---|---|---|
| 27 | 140.172 | 220.28483333333335 |
| 31 | 282.0986111111111 | 405.666 |
| 32 | 84.64241666666666 | 231.68044444444445 |
#### Chart
| Category | yolk | embryo |
|---|---|---|
| 27 | 2.044111111111111 | 2.1022777777777777 |
| 31 | 6.372111111111111 | 6.239466666666667 |
| 32 | 1.4668666666666668 | 3.1022777777777777 |
#### Chart
| Category | yolk | embryo |
|---|---|---|
| 27 | 180.64516666666665 | 6.997277777777778 |
| 31 | 309.23066666666665 | 21.995444444444445 |
| 32 | 90.6533888888889 | 65.03744444444445 |gpd1
gpd1l
gpd1c
gpd2
slc2a2
*I
*I
*S
*S
*ISP
c
b
b
a
ab
a
a
a
a
a
a
a
slc2a1b
slc2a2
slc2a5
slc2a8
slc2a11a
#### Chart
| Category | yolk | embryo |
|---|---|---|
| 27 | 4.029888888888889 | 7.530333333333334 |
| 31 | 1.5694444444444444 | 6.525533333333334 |
| 32 | 1.2307777777777777 | 1.8508333333333333 |
#### Chart
| Category | yolk | embryo |
|---|---|---|
| 27 | 180.64516666666665 | 6.997277777777778 |
| 31 | 309.23066666666665 | 21.995444444444445 |
| 32 | 90.6533888888889 | 65.03744444444445 |
#### Chart
| Category | yolk | embryo |
|---|---|---|
| 27 | 2.7583888888888883 | 1.357 |
| 31 | 2.0974444444444447 | 2.551888888888889 |
| 32 | 1.8149444444444445 | 4.765833333333334 |
#### Chart
| Category | yolk | embryo |
|---|---|---|
| 27 | 16.696733333333334 | 10.712277777777778 |
| 31 | 18.76677777777778 | 14.905777777777777 |
| 32 | 4.122 | 4.566888888888889 |
#### Chart
| Category | 卵黄 | 胚 |
|---|---|---|
| 27 | 4.04925 | 6.978833333333334 |
| 31 | 4.766166666666667 | 6.064 |
| 32 | 4.101066666666666 | 4.102533333333334 |c
b
NS
*ISP
*IS
*S
NS
c
bc
b
ab
ab
a
a
ab
b
ab
a
a
ab
a
a
a
a
a
a
a
a
a
slc2a11b
gys1
gys2
pygb
pygm
#### Chart
| Category | 卵黄 | 胚 |
|---|---|---|
| 27 | 1.4101666666666668 | 1.7305333333333333 |
| 31 | 2.0348 | 0.977388888888889 |
| 32 | 0.7170000000000001 | 0.8174444444444445 |
#### Chart
| Category | yolk | embryo |
|---|---|---|
| 27 | 2.6114444444444445 | 2.8550000000000004 |
| 31 | 2.918277777777778 | 4.738866666666667 |
| 32 | 4.393388888888889 | 5.527722222222223 |
#### Chart
| Category | yolk | embryo |
|---|---|---|
| 27 | 26.76527777777778 | 14.540500000000002 |
| 31 | 38.49316666666667 | 19.704944444444447 |
| 32 | 4.249833333333334 | 8.357833333333332 |
#### Chart
| Category | yolk | embryo |
|---|---|---|
| 27 | 29.01422222222222 | 14.709111111111111 |
| 31 | 31.205 | 31.612388888888887 |
| 32 | 7.465888888888888 | 26.715611111111112 |
#### Chart
| Category | yolk | embryo |
|---|---|---|
| 27 | 1.744666666666667 | 2.1361666666666665 |
| 31 | 1.5854444444444444 | 21.962166666666672 |
| 32 | 1.3868333333333334 | 59.799 |b
NS
*SP
*SP
NS
*ISP
b
c
ab
ab
bc
abc
a
a
a
a
ab
ab
a
a
a
a
a
Fig. S2. Changes in expression levels of the selected genes during development.
The expression levels of gluconeogenesis-related genes and glucose transporter (GLUT) gene (slc2a) family were measured. slc2a2 were measured. Horizontal and vertical axes indicate developmental stages and mRNA levels (x 10^9^ copies / g RNA), respectively. Open and filled columns indicate the mRNA levels in the yolk sac membrane and embryos, respectively. Data are presented as mean ± standard error (N = 6), and different letters indicate significant differences (P < 0.05) between groups. Tests for significance were performed by two-way ANOVA and Tukey's post-hoc test. *I indicates a significant interaction between the two factors (developmental stage and site), and *S and *P indicate significant main effects of developmental stage and the position (yolk sac or embryo), respectively.

### Slide 3
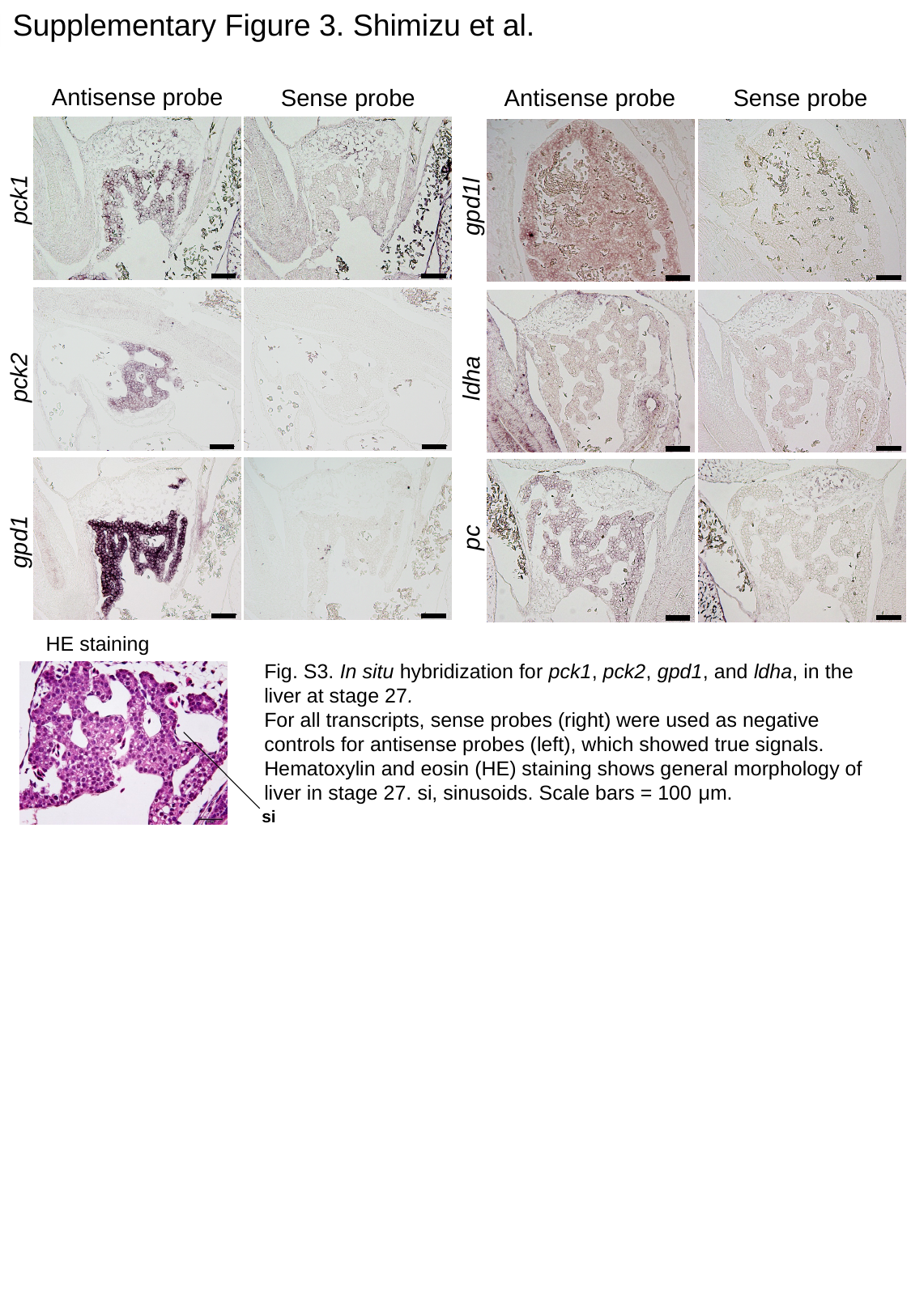

Supplementary Figure 3. Shimizu et al.
Antisense probe
Sense probe
Antisense probe
Sense probe
pck1
gpd1l
pck2
ldha
pc
gpd1
HE staining
Fig. S3. In situ hybridization for pck1, pck2, gpd1, and ldha, in the liver at stage 27.
For all transcripts, sense probes (right) were used as negative controls for antisense probes (left), which showed true signals. Hematoxylin and eosin (HE) staining shows general morphology of liver in stage 27. si, sinusoids. Scale bars = 100 μm.
si

### Slide 4
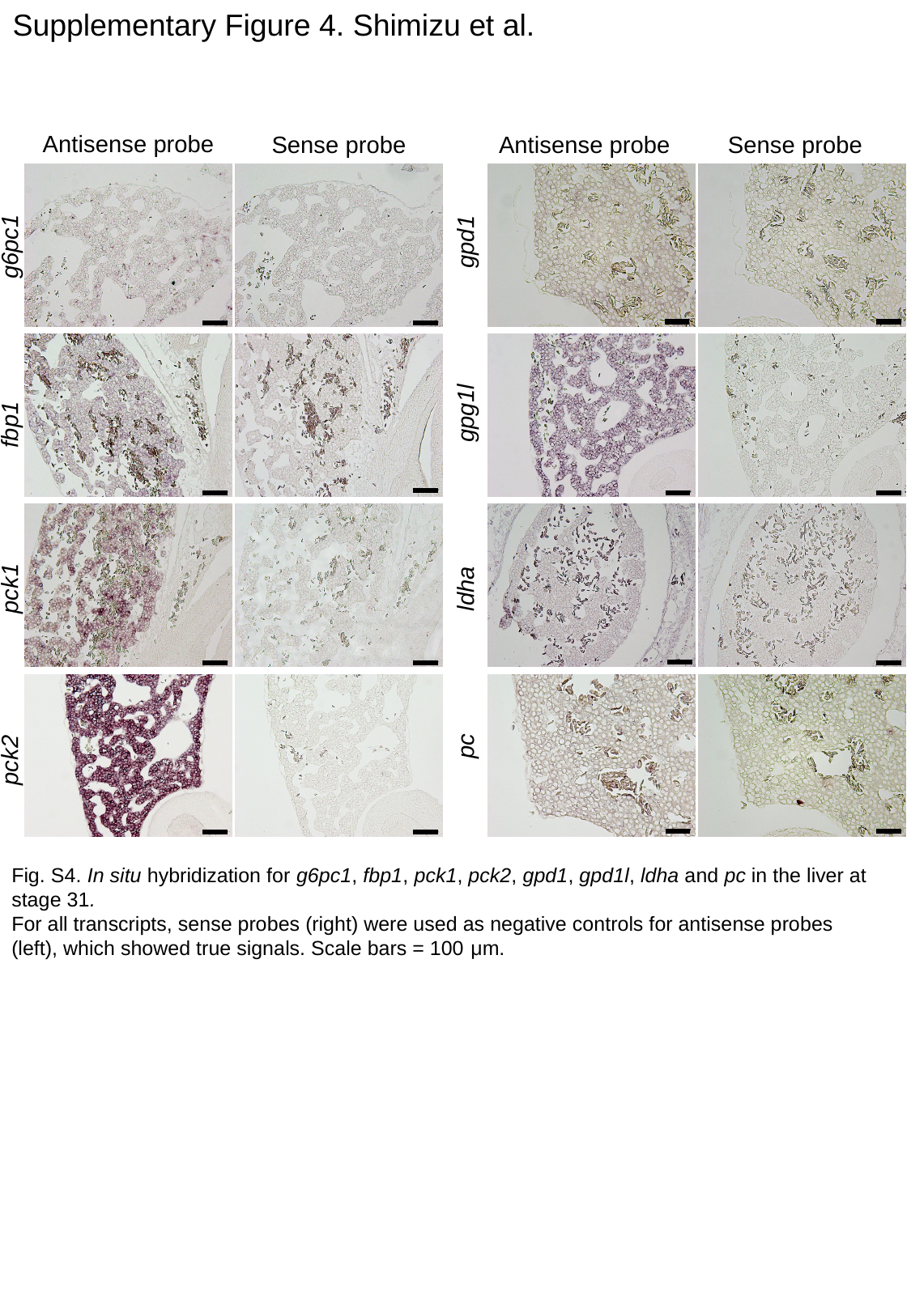

Supplementary Figure 4. Shimizu et al.
Antisense probe
Sense probe
Antisense probe
Sense probe
gpd1
g6pc1
gpg1l
fbp1
ldha
pck1
pc
pck2
Fig. S4. In situ hybridization for g6pc1, fbp1, pck1, pck2, gpd1, gpd1l, ldha and pc in the liver at stage 31.
For all transcripts, sense probes (right) were used as negative controls for antisense probes (left), which showed true signals. Scale bars = 100 μm.

### Slide 5
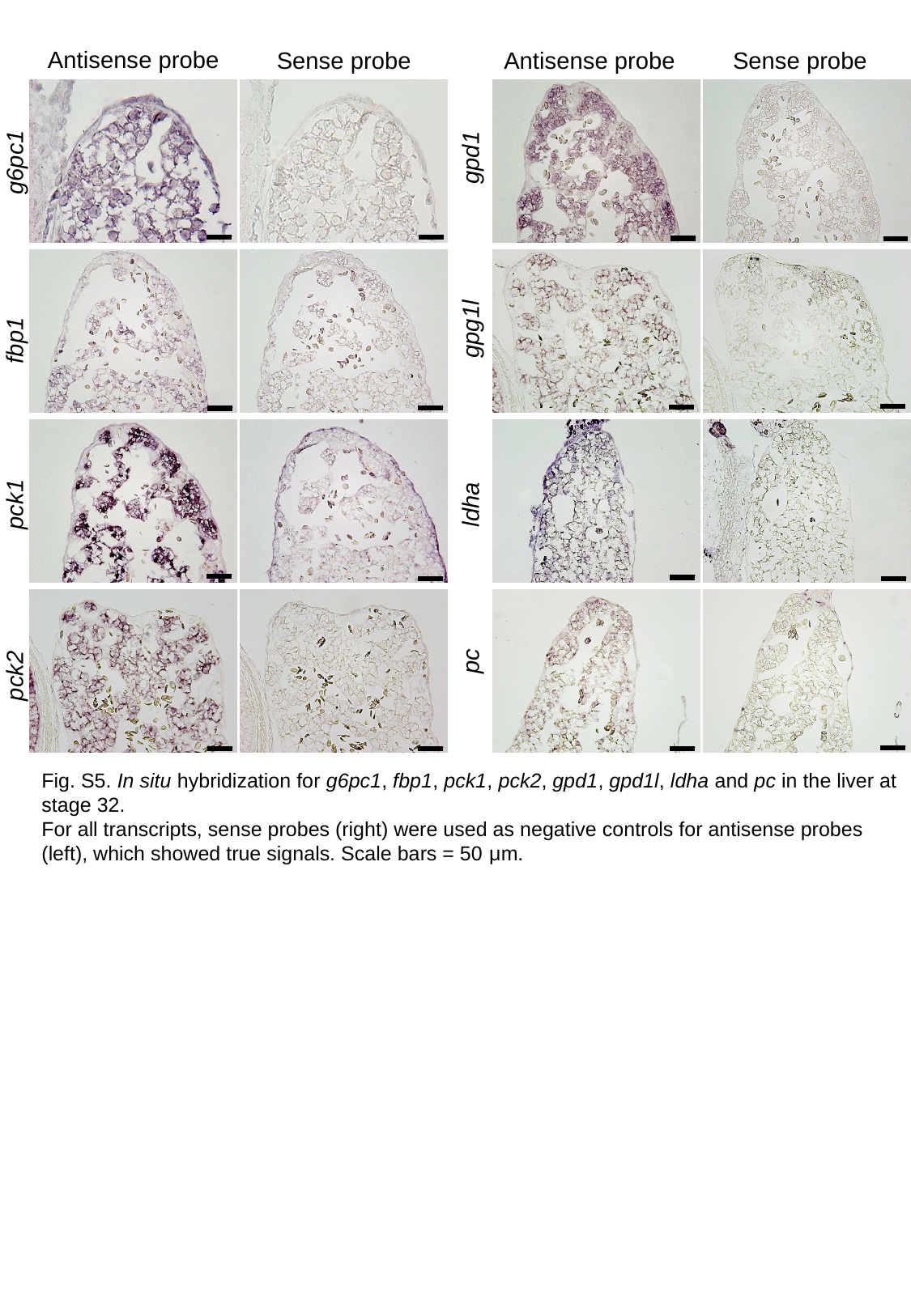

Antisense probe
Sense probe
Antisense probe
Sense probe
gpd1
g6pc1
gpg1l
fbp1
ldha
pck1
pc
pck2
Fig. S5. In situ hybridization for g6pc1, fbp1, pck1, pck2, gpd1, gpd1l, ldha and pc in the liver at stage 32.
For all transcripts, sense probes (right) were used as negative controls for antisense probes (left), which showed true signals. Scale bars = 50 μm.
